# Natural-abundance ^17^O NMR spectroscopy of magnetically aligned lipid nanodiscs

**DOI:** 10.1101/2020.05.30.125492

**Authors:** Thirupathi Ravula, Bikash Sahoo, Xiaofeng Dai, Ayyalusamy Ramamoorthy

## Abstract

Natural-abundance ^17^O NMR experiments are used to investigate the hydrated water in magnetically alligned synthetic polymer based lipid-nanodiscs. Residual quadrupole couplings (RQCs) measured from the observed five ^17^O (central and satellite) transitions, and molecular dynamics simulations, are used to probe the ordering of water moecules across the lipid bilayer.

---

Water is central to all life, and crucial for the structure and function of all biological systems. Study of hydration dynamics can provide valuable insights into the role of water in the function of such biological systems including protein, RNA and biomaterials.^1, 2^ For example, water significantly influence the structure, dynamics and the variety of functional properties of the cell membrane.^2, 3^ However, there are numerous challenges for high-resolution probing of the structure and dynamics of water molecules in a membrane environment. Herein, we demonstrate the use of natural-abundance ^17^O NMR spectroscopy to investigate the interaction of water molecules with lipid-nanodiscs.

While the very low-abundance (^~^0.037%) and a small gyromagnetic ratio (^~^1/7th of ^1^H) render ^17^O NMR to be insensitive, magic angle spinning (MAS) studies on solids well utilized the large chemical shift span and quadrupolar interaction of ^17^O.^4, 5^ ^17^O quadrupole central transition has also been used to study chemical and biological macromolecules in solution as reported in the literature.^4, 6–9^ In this study, polymer nanodiscs are used as membrane mimetics to investigate the interaction of water with lipid bilayer using natural-abundance ^17^O NMR spectroscopy. Polymer nanodiscs consists of a planar lipid bilayer encased by the polymer belt (Fig.1A).^10, 11^ Lipid nanodiscs technology is increasingly used for the structure and functional studies of membrane proteins.^12, 13^ While isotropic nanodiscs are used in solution NMR studies, the magnetic-alignment properties of polymer nanodiscs (diameter >20 nm) have been shown to be unique in enabling the application of solid-state NMR experiments to study lipids and membrane proteins, and to measure RDCs (residual dipolar couplings) using solution NMR studies.^14–17^ Natural-abundance ^17^O NMR spectra of magnetically-aligned and flipped nanodiscs are remarkable in showing all the five transitions for the spin 5/2 system, and the exciting feasibility of using the experimentally measured residual quadrupolar couplings (RQCs) to determine the orientation of the water molecules in the lipid bilayer. A combination of NMR and molecular dynamic simulation presented here reveals the molecular level ordering of water molecules across the lipid bilayer.

**Figure 1.**
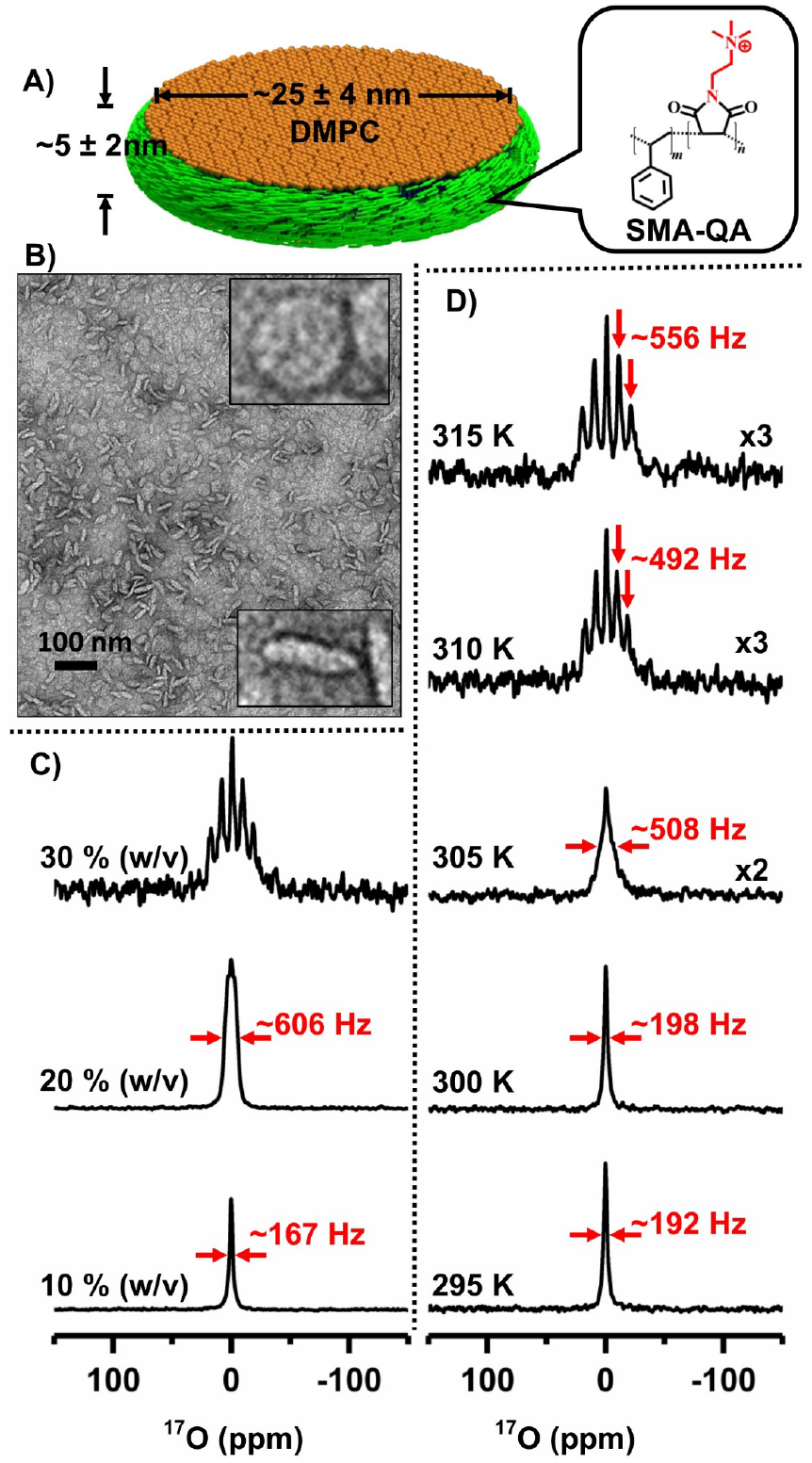
Natural-abundance ^17^O NMR spectra of polymer nanodiscs. (A) Schematic representation of SMA-QA polymer based lipid-nanodiscs. B) TEM image of SMA-QA:DMPC (0.5:1 w/w) nanodiscs. ^17^O NMR spectra of water from SMA-QA:DMPC (0.5:1 w/w) nanodiscs (^~^25 nm diameter) for varying lipid concentration at 310 K (C) and at the indicated temperatures for 30% w/v lipid concentration (D). All other details on the sample preparation, characterization, and experimental conditions are given in the supporting information.

Nanodiscs were prepared by the addition of a synthetic polymer (styrene maleimide quaternary ammonium (SMA-QA); synthesis, purification and characterization can be found elsewhere^15^) and DMPC (1,2-dimyristoyl-sn-glycero-3-phosphocholine) lipids as described in the Supporting Information, and characterized using biophysical experiments as reported previously.^18^ The polymer nanodiscs prepared in various concentrations and their ability to magnetically align in the presence of an external magnetic field were confirmed using ^31^P NMR as reported previously.^16, 18^ In addition, TEM images were used to confirm the approximate size and shape of the nanodiscs (see Figure 1B). ^17^O NMR experiments on varying concentrations of SMA-QA-DMPC nanodiscs were performed at 310 K, and the acquired spectra are shown in Figure 1(C). At a low lipid concertation (10 % w/v), a single peak corresponding to the isotropic chemical shift frequency of water was observed. The scalar coupling between ^17^O and ^1^H resulting in a triplet for isotropic water is not observed for water in the nanodiscs sample as the observed line is broader (line width is ^~^167 Hz) than the scalar coupling (^~^78 Hz). When the lipid concertation was increased to 20% w/v, the line-width of the observed ^17^O peak increased to ^~^606 Hz and also showed a triplet pattern.

Further increase in the concentration of nanodiscs to 30% w/v showed a well-resolved pentet pattern as shown in Fig.1C. Motion of water molecules in the nanodiscs sample averaged the ^17^O quadrupole coupling to a smaller value (called RQCs) such that all the five (one central and four satellite) transitions observed in the form of a pentet pattern. The RQC values are obtained by measuring the frequency difference between any of the two adjacent peaks from the pentet signal; a value of ^~^492 Hz is measured for 30% w/v nanodiscs. These results suggest that water in magnetically-aligned nanodiscs is partially aligned.

To measure the effect of temperature on the alignment of water molecules in nanodiscs, ^17^O spectra were recorded as a function of temperature (Fig.1D). As demonstrated using ^31^P and ^14^N NMR experiments, the nanodiscs align at temperatures above the main gel to liquid crystalline phase transition temperature of DMPC (T_m_ ^~^24 °C).^16^ It is remarkable that ^17^O RQC values observed for water in nanodiscs reveal the magnetic-alignment of nanodiscs as shown in Fig.1D. ^17^O spectra show an isotropic peak below T_m_ (see the spectra at 295 and 300 K with line-widths <200 Hz), the increasing temperature resulted in line-broadening near T_m_, (see the spectrum at 305 with line-width ^~^508 Hz) and a multiplet pattern is observed above T_m_ as shown for the spectrum at 310 K with an RQC of ^~^492 Hz. Further increase in the temperature to 315 K, increased the observed quadrupolar coupling to ^~^556 Hz. These observations clearly demonstrate the use of ^17^O NMR spectra to probe the water molecules ordering using magnetic-alignment of nanodiscs. To gain further insights into the dynamics of water, we measured the longitudinal relaxation (T_1_) times for all five ^17^O peaks observed (Figure 3). The T_1_ values for water molecules in the nanodiscs are in the range of 4 ms, and much smaller than that for bulk water (^~^9 ms, Fig.S2), suggesting the restricted motion of water molecules in nanodiscs under these conditions.

Since the observed RQC depends on the order parameter as well as on the orientation of bilayer-associated water molecules, a single ^17^O RQC value obtained from aligned nanodiscs is not sufficient to completely describe the alignment of water molecules. A complete description of the alignment of water molecules requires RDC values measured under different alignment conditions. To measure the RQCs of ^17^O under different alignment conditions, we used a paramagnetic salt, YbCl_3_, to flip the orientation of magnetically-aligned nanodiscs as described in our previous study.^16 31^P and ^14^N NMR experiments were used to confirm the flipping of the aligned nanodiscs upon the addition of YbCl_3_. The observed chemical shift for ^31^P (from −13.6 ± 0.5 ppm to ^~^22.7 ± 3.7 ppm upon flipping) and doubling of ^14^N RQC (from ^~^8.9 ± 0.6 kHz to ^~^16.8 ± 0.7 kHz upon flipping) confirmed the 90° flipping of nanodiscs to result in a parallel orientation of the lipid bilayer normal to the external magnetic field direction (Figure 2). ^17^O NMR spectra of water in the flipped nanodiscs showed a significant increase in the RQC value: from 492 Hz to 690 Hz upon flipping as shown in Figure 2(A and B). ^17^O NMR spectra of varying concentrations of flipped and unflipped nanodiscs are shown in Fig.S1 along with deconvoluted spectra illustrating the multiplet pattern and RQC values. While it is remarkable that the motionally averaged water RQCs are sensitive to the alignment of nanodiscs, the 90° flipping of aligned nanodiscs did not follow (3cos^2^ϕ-1) for the observed ^17^O RQCs unlike ^31^P and ^14^N spectra; where ϕ is the angle between the bilayer normal and the magnetic field axis. The observed changes for ^31^P and ^14^N are because the axis of motional averaging for lipids (collinear with the bilayer normal) is oriented perpendicular (Figure 2E) and parallel (2J) to the magnetic field axis. On the other hand, the bilayer-associated water molecules are in exchange with the free water molecules and are randomly oriented with respect to the bilayer normal. Even though the bilayer-associated water molecules prefer to orient with their symmetry (or the bisector) axes collinear with the bilayer normal as reported for zwitterionic lipids,^19^ the observed RQC values indicate that the the motional averaging of the ^17^O quadrupolar coupling is different in flipped and unflipped nanodiscs, which is more pronounced when the concentration decreases (Fig.S1). Therefore, ^17^O RQCs can be useful to probe the molecular events at the membrane-water interface.

**Figure 2.**
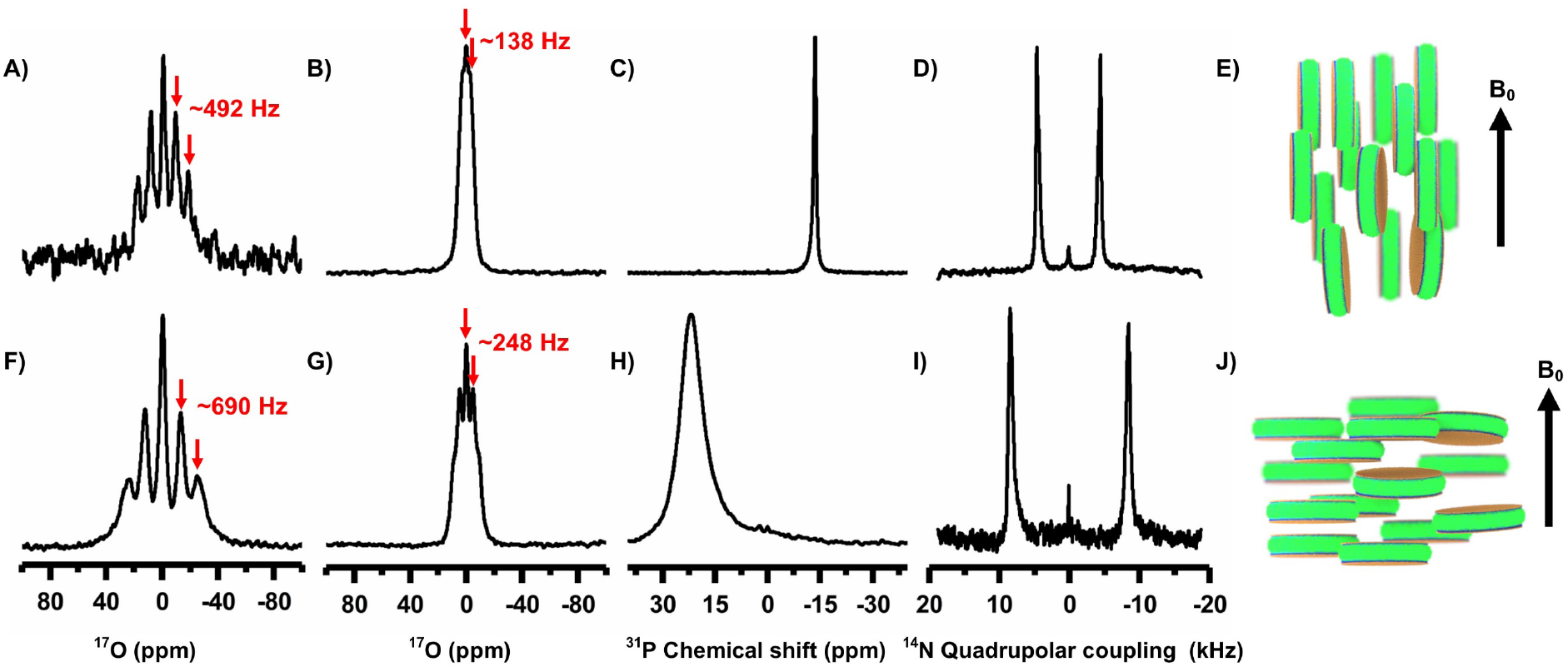
NMR spectra of magnetically-aligned and flipped nanodiscs. ^17^O (A, B, F and G), ^31^P (C and H) and ^14^N (D and I) NMR spectra of magnetically-aligned SMA-QA:DMPC nanodiscs obtained at 310 K with the lipid-bilayer-normal perpendicular (top row, as illustrated in E) and parallel (bottom row, as illustrated in J) to the external magnetic field direction. A 2 mM YbCl_3_ was used to flip the nanodiscs (bottom row). All spectra were acquired from 20% w/v lipid concentration, except that 30% w/v was used for A and F. Deconvolution of ^17^O pentet spectra of aligned and flipped nanodiscs (20% and 30% w/v) are shown in Figure S1.

**Figure 3.**
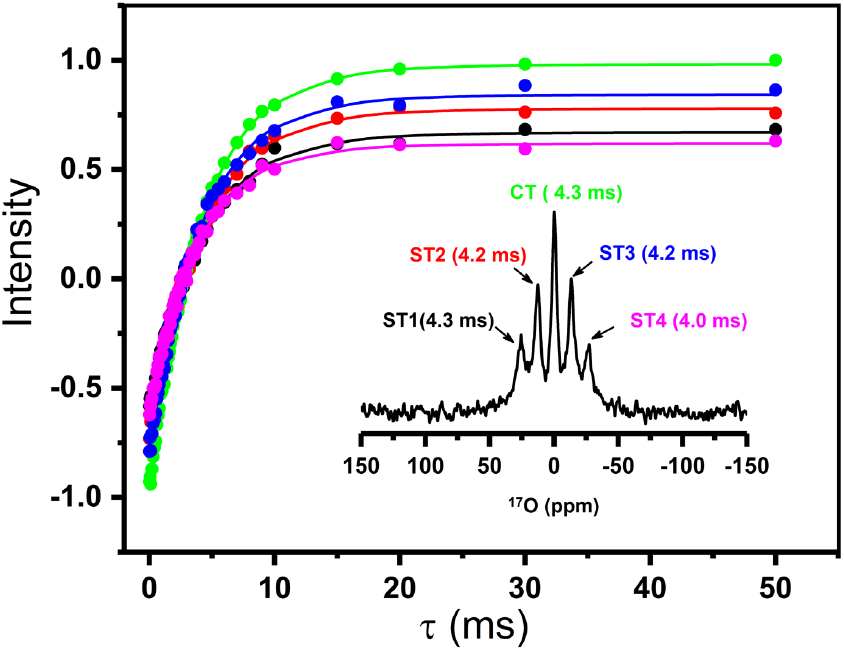
T_1_ relaxation of ^17^O NMR transitions. ^17^O spin-lattice (T_1_) relaxation for the central (CT) and satellite transitions (ST1, ST2, ST3, ST4) measured from SMAQA-DMPC (30% w/v) nanodiscs containing a 2 mM YbCl_3_. ^17^O NMR spectrum labelled with the transitions and the measured T_1_ values (inset). T_1_ values were obtained from the best-fitting of the inversion-recovery intensities (data points). See Figure S2 for ^17^O T_1_ experimental data obtained from isotropic water, and Figure S3 for ^17^O inversion recovery NMR spectra of water present in nanodiscs.

To explore the origins of an increase in the quadrupolar coupling upon 90° flipping of the nanodiscs, we used MD simulation using DMPC and water as a system (Fig.4A); additional details are given in the supporting information. The main axis of the ^17^O quadrupolar coupling tensor is perpendicular to the molecular plane of water as reported elsewhere.^20^ To obtain the relative orientations and average RQCs for the quadrupole coupling tensor across the lipid bilayer, cosθ values were calculated. MD simulations show a gradual decrease in water density across the bilayer (i.e. along the *z*-axis) and becomes zero in the hydrophobic core of the bilayer (Fig.4B). With the DMPC bilayer placed along the *xy*-plane, space and time-averaged <3cos^2^θ-1> values were calculated (Fig.4C). The resulting <3cos^2^θ-1> values are positive along the *z-*axis and negative along *x* and *y* axes. The larger <3cos^2^θ-1> magnitude along the *z*-axis suggest that the motional averaging along *x* and *y* axes more effectively reduces the RQCs than that along the *z*-axis. These results are in agreements with the observed larger RQCs for the flipped-nanodiscs, which have the bilayer-normal (i.e. the *z*-axis in Fig.4) along the magnetic field axis (Fig.2J). Thus, the measured ^17^O RQCs can be used to probe the ordering and dynamics of water molecules associated with a lipid bilayer.

**Figure 4.**
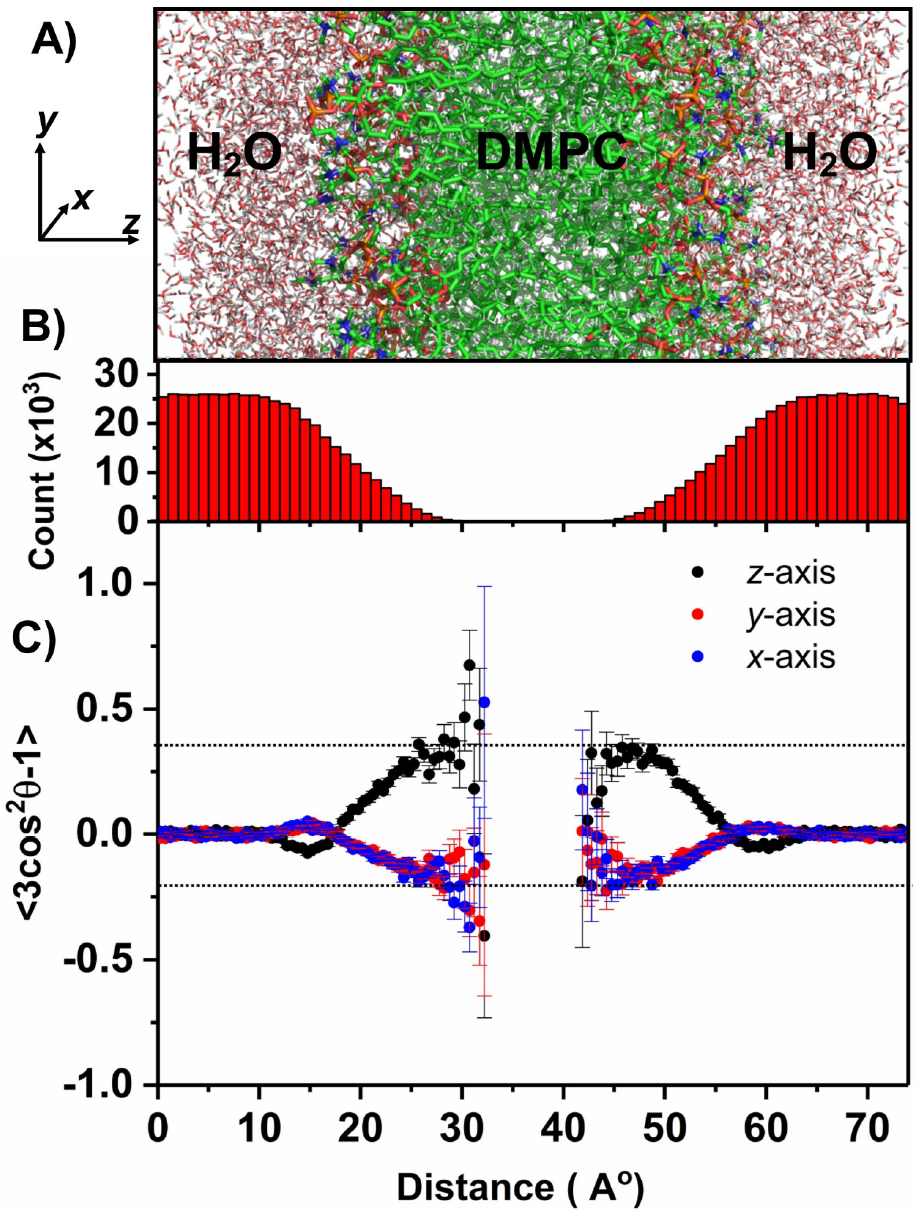
(A) Selected region of the DMPC:water system (consisting of 256 DMPC and 10153 water molecules) used in molecular dynamic simulations; additional details can be found in the supporting information (Figs.S4 and S5). *(x, y, z)* coordinates are shown with the *z*-axis parallel to the lipid bilayer normal. (B) Total number of water molecules present in the *xy* plane plotted along the z-coordinate (i.e. lipid-bilayer normal) throughout the simulation time. (C) Space and time-averaged <*3Cos^2^θ-1*> as a function of the *z*-coordinate; where θ is the angle between the normal to the water plane (i.e. the main principal axis of ^17^O quadrupole coupling tensor) and *x* (blue), *y* (red), and *z* (black) axes. Space averaging was done for all water molecules present in the *xy* plane for each point along the z-axis. The error bar represents the slandered deviation from the average value. The large error bars are due to less water molecules present in the core of the bilayer. See the supporting information for additional details on the simulations used in this study.

In conclusion, we have demonstrated that natural-abundance ^17^O NMR is an useful technique to study the magnetic-alignment of lipid-nanodiscs and also to probe membrane water interaction. At optimum hydration levels under the magnetic-alignment of nanodiscs, all five transitions of ^17^O nuclei are observed. The measured RQCs are explained based on the orientation preference of bilayer-associated water molecules. Overall, the reported results clearly indicate the molecular level ordering of water across the bilayer, and a combination of ^17^O NMR and MD simulation can be used to further probe water-membrane interactions. The reported ^17^O NMR experiments can also be used to investigate other aligned samples such as other types of nanodsics^21, 22^, bicelles^23–27^, and other alignment media used in NMR studies^28^. We also expect the reported approach and results to be useful in the investigation of the role of ordered water in many biological processes including protein folding, misfolding, amyloid aggregation, and biomineralization process.^29–31^ The use of dynamic nuclear polarization experiments^32^ to enhance ^17^O NMR sensitivity for studies on aligned samples^33, 34^ would be exciting.

## Acknowledgment

This study was supported by NIH (GM084018 to AR). We thank Dr.Ganapathy, a visiting researcher in the Ramamoorthy lab, for helping with ^17^O NMR during the initial stages of this project.

