## Supporting information for "Natural-abundance ^17^O NMR spectroscopy of magnetically aligned lipid nanodiscs"

### Materials and Methods

Poly(Styrene-co-Maleic Anhydride) cumene terminated (SMA) with a ~1.3:1 molar ratio of styrene:maleic anhydride and an average molecular weight ($M_n$) of 1600 g/mol, *N,N*-Dimethylformamide (DMF), triethylamine ($Et_3N$), HEPES, acetic acid (HOAc), hydrochloric acid (HCl), (2-aminoethyl)trimethylammonium chloride hydrochloride, and sodium hydroxide (NaOH) were purchased from Sigma-Aldrich®. 1,2-dimyristoyl-*sn*-glycero-3-phosphocholine (DMPC) was purchased from Avanti Lipids Polar, Inc®.

**Synthesis and characterization of SMA-QA polymer:** SMA-QA polymer was synthesized and characterized, and its nanodisc-forming capability was tested as reported previously.[1]

#### NMR sample preparation:

Lipid-nanodiscs were prepared using a stock solution of DMPC (10 mg/mL) and SMA-QA (10 mg/mL) in 10mM HEPES buffer containing 100 mM NaCl at pH 7.4. Nanodiscs were formed by the addition of 5 mL of SMA-QA stock solution to 10 ml of DMPC. The DMPC to SMA-QA weight ratio was 1:0.5 w/w. The resulting solution was subjected to freeze-thaw cycles between liquid nitrogen temperature and 35 °C at least 4 times. Nanodiscs samples were incubated for 12 hr at room temperature. The sample was concentrated to 1 mL using an Amicon Filter with 50-kDa membrane. The resulting nanodiscs solution was diluted to 10 mL using HEPES buffer and concentrated to 1 mL. This step was repeated 5 times to completely remove any non-nanodisc-forming (or free or aggregated) polymer from the solution. In the final washing step, the solution was concentrated to give the required lipid concentration. The resulting nanodiscs were carefully transferred to a 4 mm glass tube for NMR experiments.

#### Natural-abundance $^{17}$O NMR experiments:

NMR experiments were performed on a Bruker NMR spectrometer operating at a resonance frequency of 400.11 MHz for proton and 54.24 MHz for $^{17}$O nuclei. A 5 mm triple-resonance HXY MAS NMR probe was used under static conditions. $^{17}$O NMR spectra were acquired using a Hahn echo pulse sequence with 12 µs 90° pulse, 20 µs echo delay, 0.2 s relaxation/recycle delay, 400 ppm spectral width and 0.2 s acquisition time. Depending on the sample amount used in the experiment, the number of scans used to acquire the reported $^{17}$O NMR spectra were varied between 10,000-20,000 to obtain a reasonable signal-to-noise ratio. The acquired spectra were processed with a 50 Hz line-broadening, and the chemical shift was referenced externally using the $^{17}$O signal from water at 0 ppm. $T_1$ experiments were performed using the inversion-recovery method and using the Hahn echo acquisition.

### $^{31}$P NMR experiments:

$^{31}$P NMR experiments were performed on a Bruker NMR spectrometer operating at a resonance frequency of 400.11 MHz for proton and 161.97 MHz for $^{31}$P nuclei. A 5 mm triple-resonance HXY MAS NMR probe was used under static conditions. $^{31}$P NMR spectra were acquired using a 5 μs 90° pulse followed by a 25 kHz TPPM (two pulse phase modulation)[2] proton decoupling. 512 scans were acquired for each sample with a relaxation/recycle delay of 2.0 s.

### $^{14}$N NMR Experiments:

Nitrogen-14 NMR spectra were acquired using a Bruker 400 MHz solid-state NMR spectrometer and a 5 mm double-resonance probe operating at the $^{14}$N resonance frequency of 28.910 MHz. $^{14}$N NMR spectra were recorded using the quadrupole-echo pulse sequence[3] with a 90° pulse length of 8 μs and an echo-delay of 80 μs. $^{14}$N signal was acquired using 25 ms acquisition time, 20,000 scans and a recycle delay of 0.2 s with no $^{1}$H decoupling.

### **Molecular dynamics simulations**:

All-atom MD simulation was performed with Gromacs v5.0.7.[4] The initial coordinates of DMPC lipid-bilayer containing 256 lipids with an average surface area of 64.1 Å$^2$ and 10153 water molecules were generated using CHARMM-GUI.[5] Water thickness of 20 Å was maintained on both sides of the lipid-bilayer for MD simulation. The MD system was neutralized using 0.1 M NaCl and subjected to MD calculation by applying periodic boundary conditions. CHARMM36 force field[6] was used for DMPC lipids and TIP3P model for water molecules. The linear constraint solver (LINCS) algorithm[7] was used to constrain all bonds to their equilibrium lengths. A cutoff radius of 12 Å was used to calculate the non-bonded interactions for both short-range electrostatic and van der Walls interactions. The particle mesh Ewald method was applied to calculate the long-range interactions. The MD system was energy minimized using a steepest descent algorithm. Energy minimized systems were equilibrated using the constant particle, pressure and temperature (NPT) ensemble at 310.15 K prior to the final MD simulation. Semi-isotropic Parrinello-Rahman pressure coupling algorithm with 1 bar pressure was used to keep the surface tension constant. A final MD simulation of 100 ns with the time step set to 2 fs was performed at 310.15 K, which is above the DMPC phase transition temperature. Nose-Hoover thermostat and Parrinello-Rahman pressure coupling were used for the final MD simulation. The MD trajectory and MD snapshots were analyzed using visual molecular dynamics (VMD).[8] The space and time averaged $<3\cos^2\theta-1>$ were calculated using an in-house Perl script as shown below.

### PERL script used to calculate the space and time averaged $<3\cos^2\theta-1>$ values from nanodiscs

```
#!/usr/bin/perl -w
```

```
use strict;
use Math::Complex;
use Math::Trig;
open (FILEW, ">>out6.txt");
my @files = glob("/mnt/d/DMC/DMPC/*");
foreach my $file(@files){
open (FILER, "$file");

my @lines = <FILER>;
my $l;
my ($Ox, $Oy, $Oz);
my ($H1x, $H1y, $H1z);
my ($H2x, $H2y, $H2z);
my ($a,$b, $c);
my ($a1,$b1, $c1);
my ($a2,$b2, $c2);
my ($x, $y, $z);
my ($d1,$d2,$d3,$angle);
my ($i,$size);
$size = @lines;
for($i=1;$i<$size;$i++){
$l = $lines[$i];
        if (substr($l,0,4) eq "ATOM" && substr($l,13,3) eq "OH2") {
            $Ox = sprintf(substr($l,31,7));
            $Oy = sprintf(substr($l,39,7));
            $Oz = sprintf(substr($l,47,7));
         }
       if (substr($l,13,2) eq "H1"){
        $H1x = sprintf(substr($l, 31,7));
        $H1y = sprintf(substr($l, 39,7));
        $H1z = sprintf(substr($l, 47,7));
     }
        if (substr($l,13,2) eq "H2"){
        $H2x = sprintf(substr($l, 31,7));
        $H2y = sprintf(substr($l, 39,7));
        $H2z = sprintf(substr($l, 47,7));



        #calculation starts here
        $a1 = $H1x-$Ox;
        $b1 = $H1y-$Oy;
        $c1 = $H1z-$Oz;
        $a2 = $H2x-$Ox;
        $b2 = $H2y-$Oy;
        $c2 = $H2z-$Oz;
        my $d1 = sqrt($a1*$a1+$b1*$b1+$c1*$c1);
        my $d2 = sqrt($a2*$a2+$b2*$b2+$c2*$c2);
        $a = $b1*$c2-$b2*$c1;
        $b = $a2*$c1-$a1*$c2;
        $c = $a1*$b2-$b1*$a2;
```

```
                    my $d = sqrt($a*$a+$b*$b+$c*$c);
          $x= $a/$d;
          $y= $b/$d;
         $z= $c/$d;
                      my $z11 = 3*$z*$z-1;
                      my $y11 = 3*$y*$y- 1;
                      my $x11 = 3*$x*$x-1;

       print FILEW " ".$Oz."  ".$z11." ".$x11." ".$y11."  '\n";
        }
      }
  print "[$file]\n";

}
close FILEW;
close FILER;
```

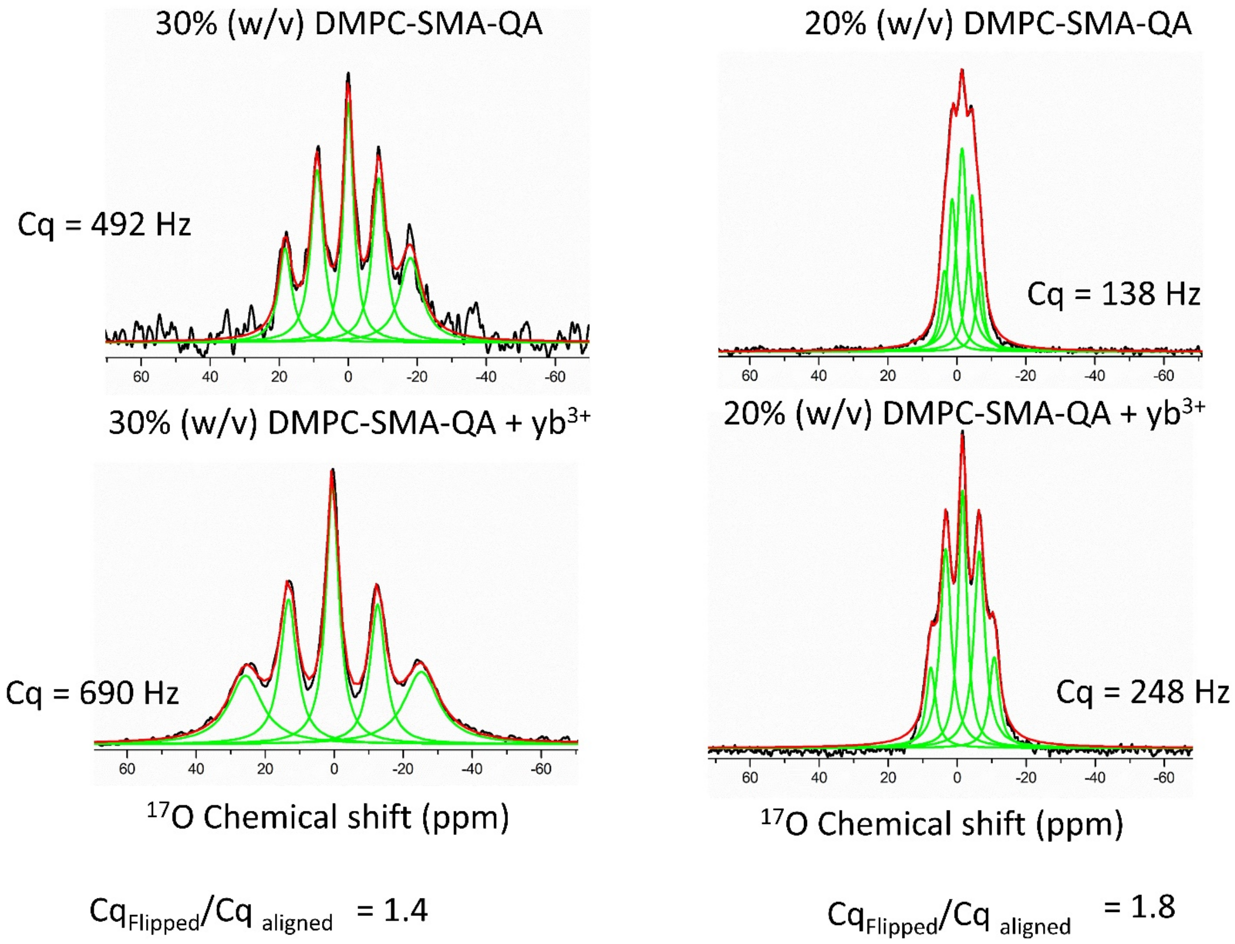


**Figure S1. Experimental measurement of $^{17}$O quadrupolar couplings.** Residual $^{17}$O quadrupolar couplings (RQC) were measured from natural-abundance $^{17}$O NMR spectra of water present in the SMA-QA:DMPC nanodiscs samples (containing 30% w/v and 20% w/v concentration of lipids) at 310 K for both aligned and flipped orientations. All NMR spectra were deconvoluted using the *Origin Lab software* and the $^{17}$O RQC (denoted as Cq) values were obtained by measuring the frequency difference between the two adjacent peaks.

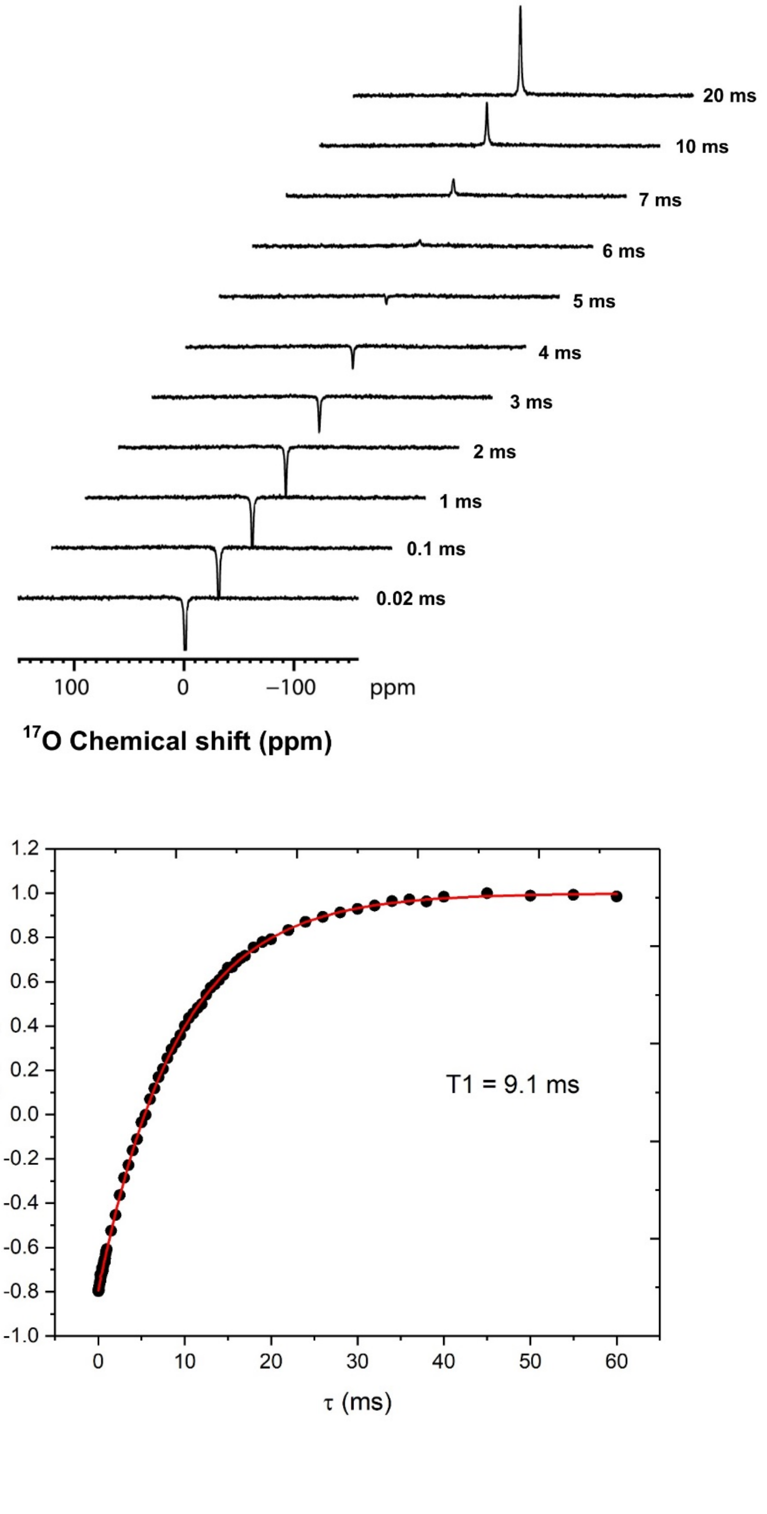


**Figure S2. Measurement of $^{17}O$ $T_1$ values**. Natural-abundance $^{17}O$ inversion-recovery NMR experiments were used to measure the $T_1$ (spin-lattice or longitudinal) relaxation for a water sample containing 10 mM HEPES and 100 mM NaCl at 310 K. Selective spectral traces are shown at the top. The best-fit (red) of the experimentally measured inversion recovery intensities (black) resulted in a $T_1$ value of 9.1 ms (bottom). Spin-spin (or *J* or scalar) coupling between $^1H$

and $^{17}O$ for the buffer sample is not observed due to the large linewidth; full-width at half maximum is ~ 90 Hz. On the other hand, previous studies using acetone to dilute water reported a value of 82±1 Hz for the scalar coupling between $^{1}H$ and $^{17}O$ nuclei.[9] A recent study on endofullerenes (with a single $^{17}O$-labeled water molecule trapped in each endofullerene) dissolved in nematic liquid crystal N-(4-methoxybenzylidene)-4-butylaniline (MBBA) reported a pentet multiplet for $^{17}O$ with a splitting of 10.52 kHz, and each of the five lines is a 1:2:1 triplet with a splitting of 208 Hz due to the combination of -78 Hz O-H scalar coupling and residual $^{17}O$-$^{1}H$ dipolar coupling.[10]

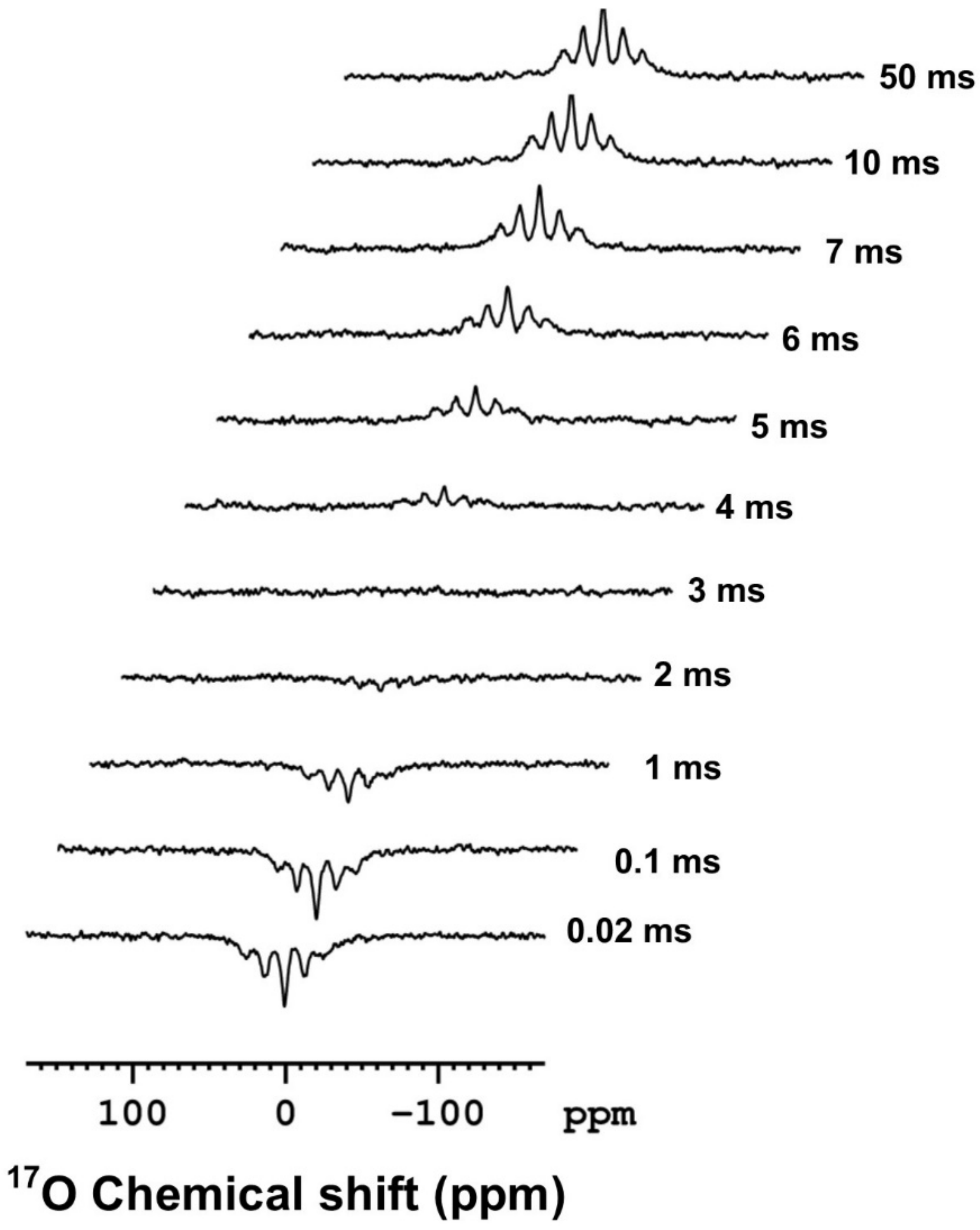


**Figure S3. $T_1$ values measured for $^{17}O$.** Natural-abundance oxygen-17 inversion recovery experimental spectra of SMAQA-DMPC (30% w/v) nanodiscs containing 2 mM $YbCl_3$ used to measure the spin-lattice (or longitudinal) relaxation time, $T_1$. Only selected spectral traces are shown here. The $T_1$ values for the main and satellite $^{17}O$ transitions are shown in Figure 3 in the main text.

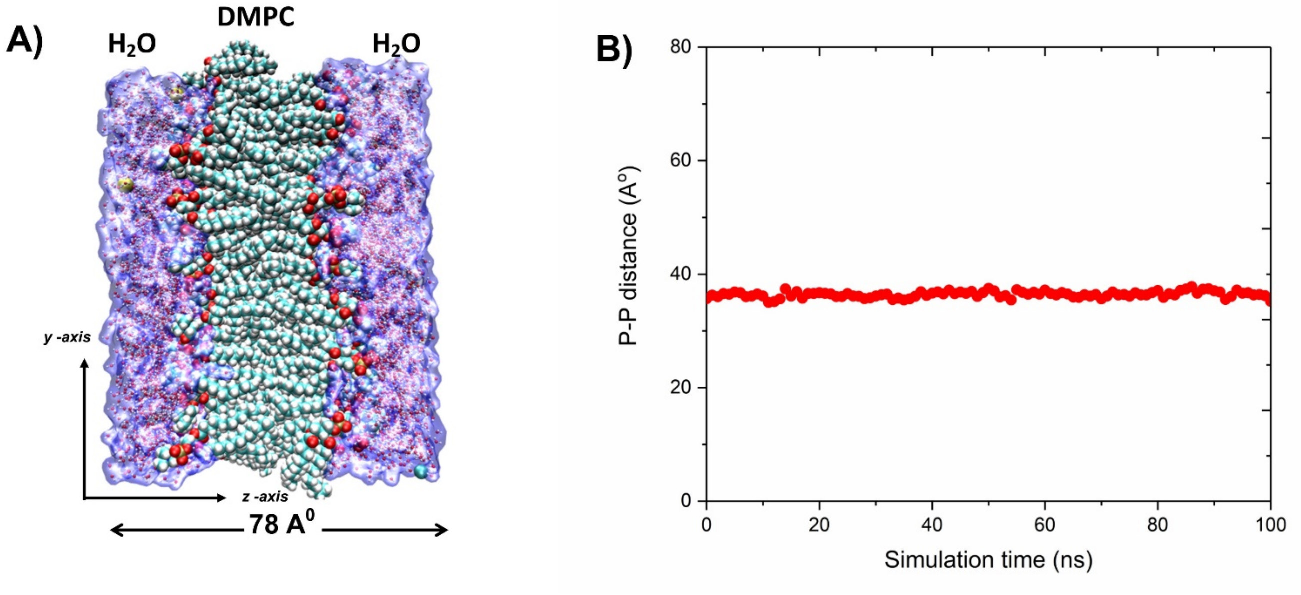


**Figure S4. Molecular dynamics simulations used to calculate <3cos$^2$θ-1> values.** (A) DMPC:water system used for the MD simulation and (B) average phosphorus - phosphorus distance as function of the simulation time.

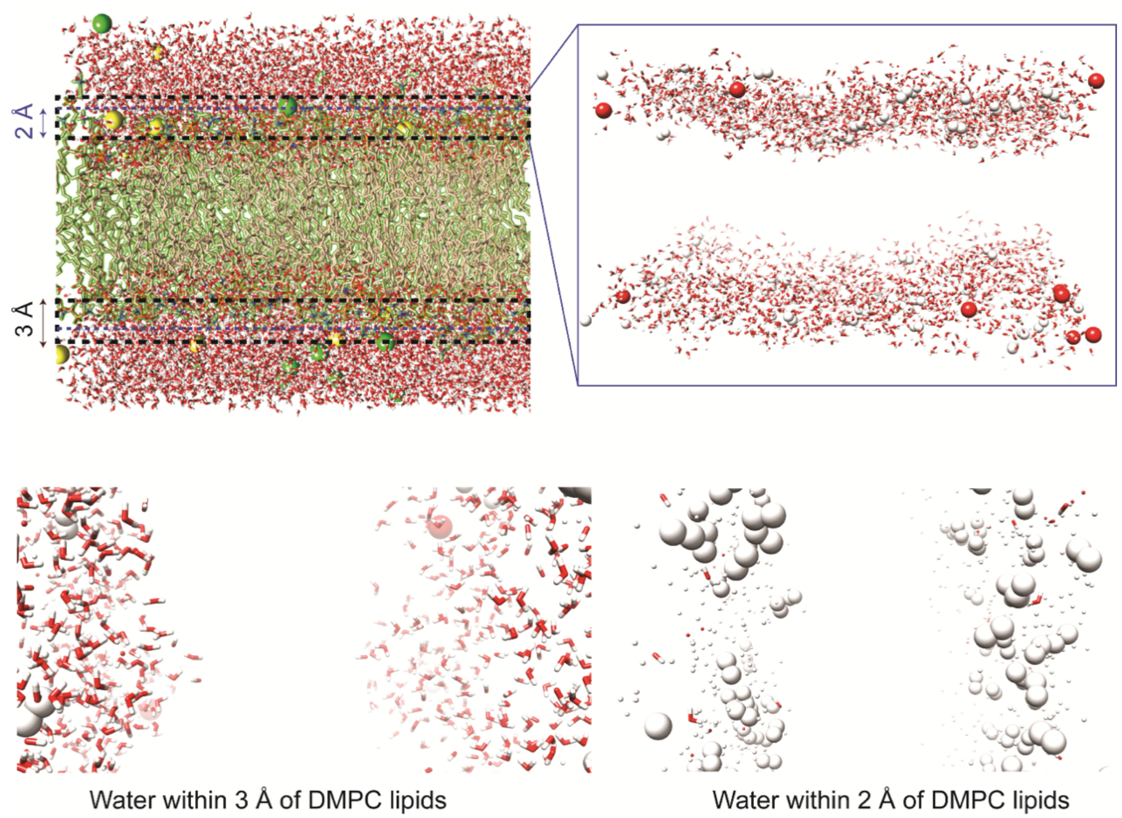


**Figure S5. Simulated DMPC-water system.** MD snapshot retrieved from 100 ns MD simulations showing the orientation of water molecules distributed within 2 Å and 3 Å from DMPC lipids, as indicated by dashed lines. (Bottom) Zoom in images (90° rotation with respect to the image at the top) for the water molecules as indicated.
